# Type III CRISPR-Cas systems generate cyclic oligoadenylate second messengers to activate Csm6 RNases

**DOI:** 10.1101/153262

**Authors:** Ole Niewoehner, Carmela Garcia-Doval, Jakob T. Rostøl, Christian Berk, Frank Schwede, Laurent Bigler, Jonathan Hall, Luciano A. Marraffini, Martin Jinek

## Abstract

In many prokaryotes, type III CRISPR–Cas systems detect and degrade invasive genetic elements by an RNA-guided, RNA-targeting multisubunit interference complex that possesses dual RNase and DNase activities. The CRISPR-associated protein Csm6 additionally contributes to interference by functioning as a standalone ribonuclease that degrades invader RNA transcripts, but the mechanism linking invader sensing to Csm6 activity is not understood. Here we show that Csm6 proteins are activated through a second messenger generated by the type III interference complex. Upon target RNA binding by the type III interference complex, the Cas10 subunit converts ATP into a cyclic oligoadenylate product, which allosterically activates Csm6 by binding to its CARF domain. CARF domain mutations that abolish allosteric activation inhibit Csm6 activity *in vivo*, and mutations in the Cas10 Palm domain phenocopy loss of Csm6. Together, these results point to a hitherto unprecedented mechanism for regulation of CRISPR interference that bears striking conceptual similarity to oligoadenylate signalling in mammalian innate immunity.

---

Clustered regularly interspaced short palindromic repeat (CRISPR) loci and CRISPR-associated (Cas) proteins constitute an adaptive prokaryotic immune system directed against invasive genetic elements such bacteriophages and plasmids^1-3^. Class 1 CRISPR-Cas systems, comprising type I, III and IV systems, mediate nucleic acid interference by means of a multiprotein-RNA complex composed of a processed CRISPR RNA (crRNA) and several Cas protein subunits^4^. In type III systems, the interference complex (known as Csm complex in type III-A/D systems and Cmr complex in type III-B/C systems) is assembled from the signature multidomain protein Cas10 (Csm1 in type III-A/D or Cmr2 in type III-B/C systems) and additional Cas proteins (Csm2-5 in type III-A/D, or Cmr1 and Cmr3-6 in type III-B/C)^5^. In these systems, interference is transcription-dependent and is thought to be initiated by the crRNA-guided interference complex binding to the nascent transcript of the target gene^6-8^. The target RNA-bound complex functions as a sequence-specific endoribonuclease (RNase) to cleave the bound target RNA and in addition has a target RNA-stimulated non-specific deoxyribonuclease (DNase) activity that cleaves single-stranded DNA^6,9-12^.

Members of the Csm6 or the related Csx1 protein families are frequently encoded within type III CRISPR-Cas systems^13^. These proteins are characterized by the presence of an N-terminal CRISPR-Associated Rossmann Fold (CARF) domain and a C-terminal Higher Eukaryotes and Prokaryotes Nucleotide-binding (HEPN) RNase domain^13,14^. In the *Staphylococcus epidermidis* type III system, Csm6 is required for efficient antiplasmid and antiviral immunity^15,16^. Csm6 proteins function as RNases^16-18^ that have been shown to nonspecifically degrade invader-derived RNA transcripts, providing an additional interference mechanism that complements the nuclease activities of the type III interference complex^16^. As Csm6 ribonucleases do not appear to physically associate with the Cas10-containing interference complex^19-21^, the mechanism linking their nuclease activity to invader recognition is not known.

## Allosteric activation of CRISPR-associated Csm6 ribonucleases

We previously showed that Csm6 is a homodimeric RNase that contains a conserved active site at the dimer interface of the HEPN domains^17^. Our structural studies further revealed that Csm6 contains a conserved, positively charged cleft at the dimeric interface of the CARF domains. Given the dimeric architecture of Csm6 and the fact that the CARF domain is a variant of the Rossmann fold, which is typically found in nucleotide-binding proteins^13^, we hypothesized that the ribonuclease activity of Csm6 might be allosterically regulated by a twofold symmetric nucleotide ligand. To address this, we initially tested the ribonuclease activity of *Thermus thermophilus* Csm6 (TtCsm6) in the presence of cyclic bis-(3’-5’)-adenylate (c-di-AMP), a well-known bacterial second messenger^22^. We observed that TtCsm6 activity was substantially increased in the presence of c-di-AMP (Extended Data Fig. 1). However, the stimulatory effect was batch-dependent; c-di-AMP batches prepared by chemical synthesis were active, while enzymatically-generated c-di-AMP had no effect (Extended Data Fig. 1), suggesting that activation was due to a contaminating by-product of chemical synthesis. Because chemical synthesis of cyclic dinucleotides typically involves cyclization of a linear dinucleotide precursor^23^, we speculated that the true activator might be a linear or cyclic oligoadenylate nucleotide generated by dinucleotide concatenation. To test this hypothesis, we first analysed the activation of TtCsm6 by linear tetraadenylates in ribonuclease activity assays that employed either a fluorophore-labelled ssRNA substrate (Fig. 1a) or a fluorogenic RNA substrate containing a fluorophore and a quencher whose cleavage results in a quantifiable increase in fluorescence signal (Fig. 1b, Extended Data Fig. 2). TtCsm6 was potently activated by tetraadenylate (A4); moreover, the presence of a 2’,3’-cyclic phosphate group further potentiated activation (Fig. 1a, b). Profiling a panel of oligonucleotides of varying lengths revealed that the 2’,3’-cyclic phosphate-terminated tetradenylate (A4>P) had the strongest effect on TtCsm6 (Fig. 1c), although longer oligoadenylates (oligoA) were also modestly active (Fig 1c, Extended Data Fig. 2b). To test whether oligoA-mediated activation was a general property of Csm6 enzymes, we additionally examined evolutionarily divergent Csm6 proteins from *Methanothermobacter thermoautotrophicus* (MtCsm6, type III-D) and *Enterococcus italicus* (EiCsm6, type III-A) (Extended Data Fig. 2a). Both MtCsm6 and EiCsm6 were efficiently activated by oligoA nucleotides (Extended Data Fig. 2c, d), with MtCsm6 displaying strongest activation by A4>P (Extended Data Fig. 3) and EiCsm6 by 2’,3’-cyclic phosphate-terminated hexaadenylate (A6>P, Fig. 1d). Together, these results demonstrate that Csm6-family ribonucleases are allosterically activated by oligoadenylate nucleotides.

**Figure 1.**
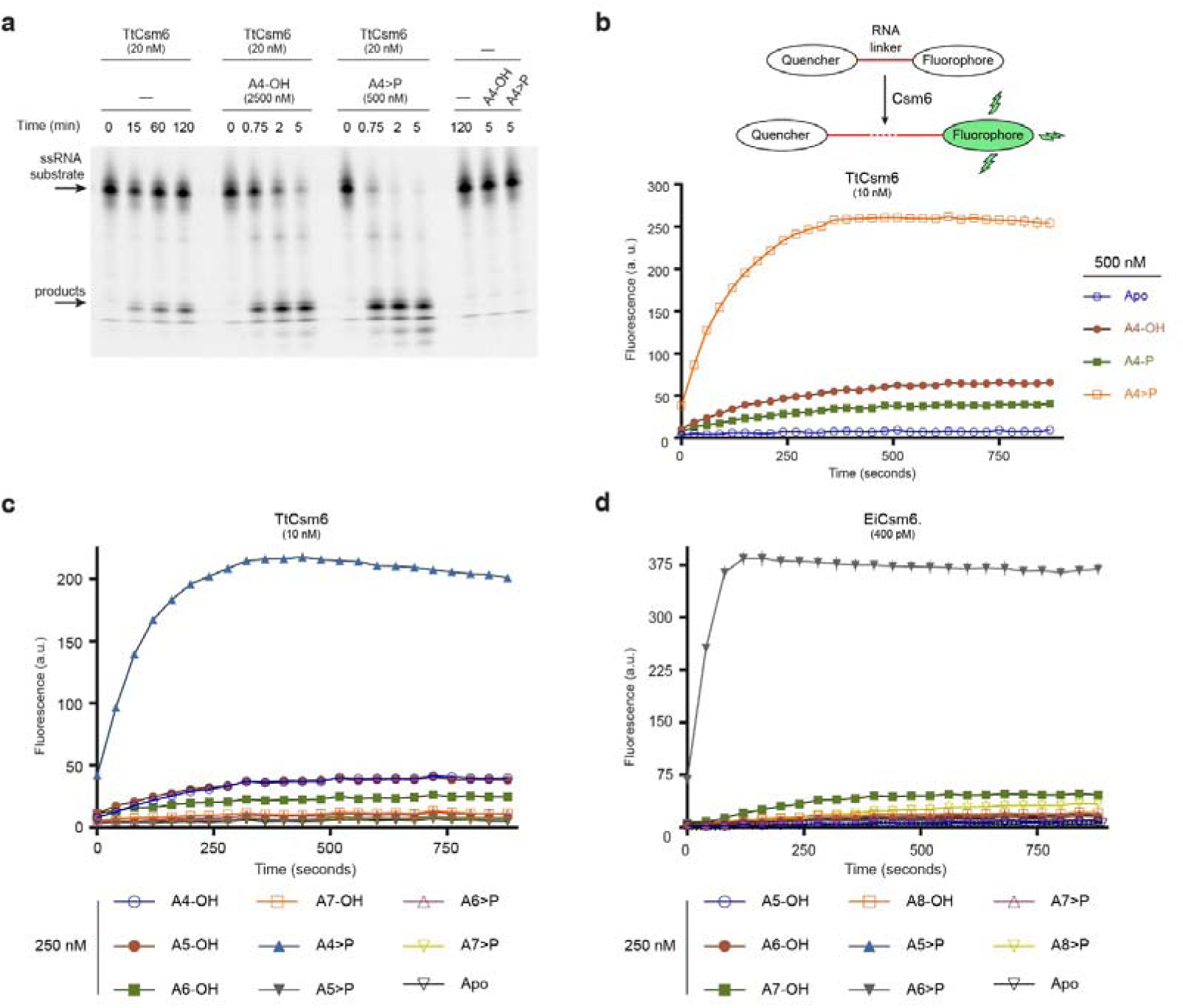
Csm6 is allosterically activated by oligoA nucleotides. **a,** TtCsm6 ribonuclease activity assay using a Cy5-labelled ssRNA substrate, in the presence of linear tetraadenylate nucleotides containing 3’-hydroxyl (A4-OH) or 2’,3’-cyclic phosphate (A4>P) groups. **b**, Top: schematic representation of fluorogenic ribonuclease activity assay; bottom: TtCsm6 RNase activity in the presence of tetraadenylates containing 3’-OH (A4-OH), 3’-phosphate (A4-P) or 2’,3’-cyclic phosphate (A4>P) groups. **c**, TtCsm6 RNase activity in the presence of oligoadenylates of varying lengths containing 3’-OH or 2’,3’-cyclic phosphate. **d**, EiCsm6 RNase activity in the presence of oligoadenylates of varying lengths containing 3’-OH or 2’,3’- cyclic phosphate. All data points represent the mean of three replicates ± s.e.m.

## The CARF domain of Csm6 is an oligoA sensor

To quantify the activation of Csm6 ribonucleases, we analysed the ribonuclease activity of EiCsm6 as a function of A6>P concentration (Fig. 2a). Plotting initial reaction velocity against effector concentration and fitting the curve by non-linear regression analysis using a log(dose)-versus-response relationship gave a half-maximal effective concentration (EC50) of ~60 nM (Fig. 2b). Since HEPN domain ribonucleases typically bind their RNA substrates with micromolar affinities^24,25^, this strongly suggests that the A6>P effector is recognized by an allosteric binding site in EiCsm6 that is distinct from the ribonuclease active site in the HEPN domain. Based on the proposed function of CARF domains in binding nucleotide ligands^13^, we hypothesized that the CARF domain functions as the allosteric sensor of oligoA. To test this, we engineered a glutamine-to-alanine substitution in the conserved CARF domain motif Ser112–Gln116 in EiCsm6 (Extended Data Fig. 4a), which maps to the dimeric CARF domain interface in TtCsm6 and would correspond to the ligand-binding face in a canonical Rossmann fold domain (Extended Data Fig. 4b). In contrast to wild-type EiCsm6, the Q116A EiCsm6 mutant (dEiCsm6^CARF^) no longer responded to A6>P, indicating that disruption of the CARF domain motif abrogated allosteric activation (Fig. 2c,d). This result implies that the Csm6 CARF domain functions as the allosteric sensor of oligoA ligands and suggests that effector binding at the CARF domain dimer interface is transduced to the ribonuclease active site at the dimeric interface of the HEPN domains.

**Figure 2.**
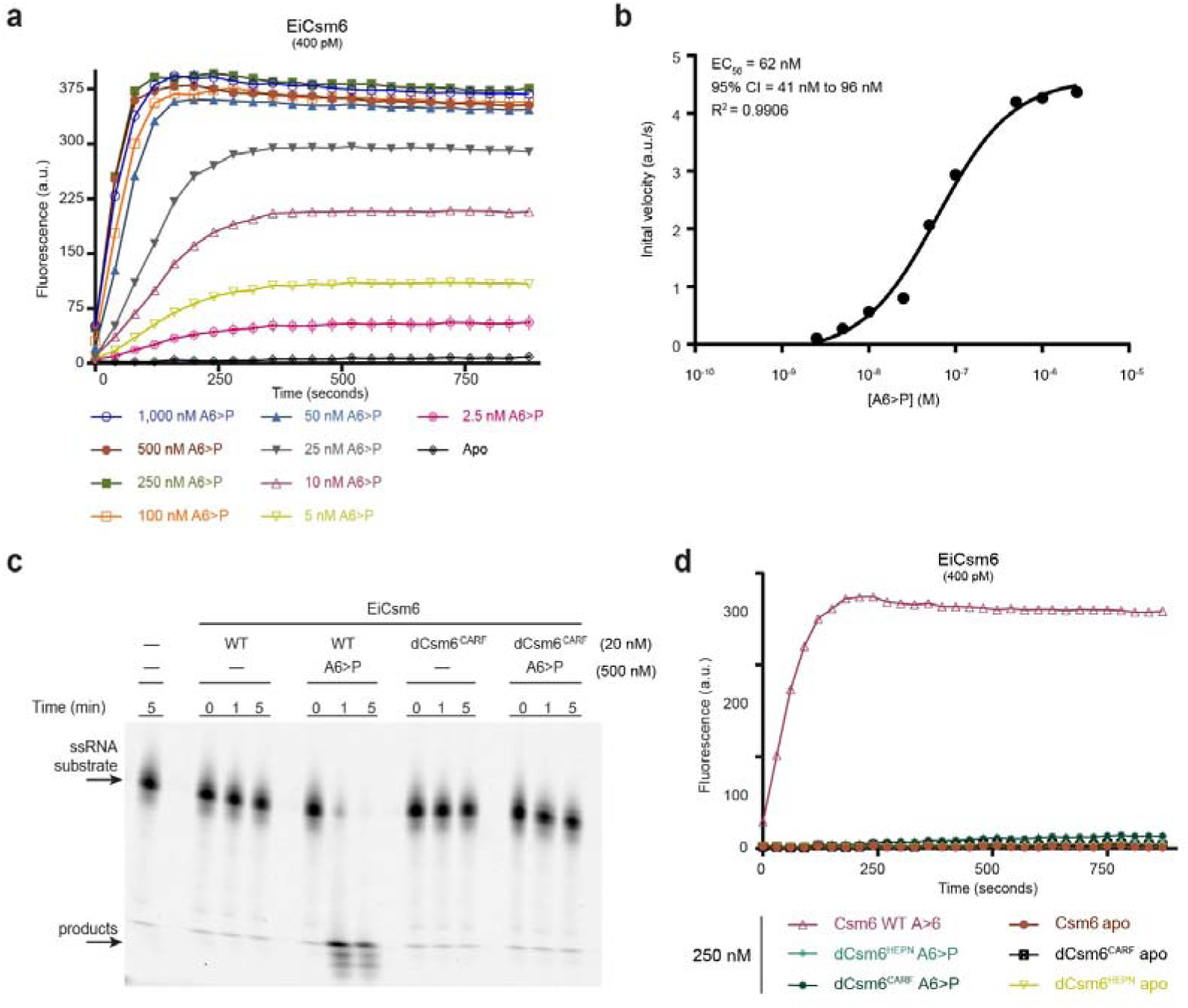
High-affinity recognition of oligoA is mediated by the Csm6 CARF domain. **a,** EiCsm6 RNase assay in the presence of varying concentrations of A6>P. **b**, Log(dose)- versus-response curve and EC50 derived from assay in panel **a**. Error in the EC_50_ value is indicated as 95% confidence interval (CI). **c**, Ribonuclease activity assay using a Cy5-labelled ssRNA and either wild type (WT) EiCsm6 or dEiCsm6^CARF^ in the presence or absence of A6>P. **d**, Ribonuclease activity assay using WT EiCsm6, dEiCsm6^CARF^ and dEiCsm6^HEPN^ proteins in the presence of A6>P. All data points represent the mean of three replicates ± s.e.m.

## Csm6 is activated by a second messenger generated by the type III interference complex

Since Csm6 proteins function in the context of type III CRISPR-Cas systems, we speculated that allosteric activation of Csm6 is dependent on an enzymatic activity harboured by the multisubunit type III CRISPR interference complex. Guided by a processed crRNA, the Cas10-containing complex binds to a target RNA, which leads to sequence-specific target RNA cleavage and concurrently stimulates sequence non-specific DNase activity of the complex^9-12^. Given that Csm6 proteins have not been found to be physically associated with the interference complex, we hypothesized that the complex would additionally generate a diffusible allosteric activator of Csm6 in a target RNA-dependent manner. We exploited the *Enterococcus italicus* type III-A system and co-expressed CRISPR-associated genes *csm1-csm6* and *cas6* together with two repeat-spacer units of the endogenous CRISPR array in *Escherichia coli* from a single polycistronic mRNA (Extended Data Fig. 5a). The purified type III-A interference complex, hereafter referred to as EiCsm(1-5), displayed subunit stoichiometry consistent with the general molecular architecture of type III interference complexes (Extended Data Fig. 5b). Moreover, the complex was capable of cleaving a cognate RNA target, yielding a characteristic pattern of cleavage products at six-nucleotide intervals, indicative of crRNA-guided ribonuclease activity mediated by the Csm3 subunits within the complex^20,26^. As expected, target RNA cleavage was dependent on the presence of a conserved aspartate residue in the Csm3 subunits (Asp32), as alanine substitution of the Csm3 aspartate (dCsm3^D32A^) resulted in loss of RNase activity (Extended Data Fig. 5c).

To test whether the allosteric effector of Csm6 is produced by the type III interference complex, we incubated the EiCsm(1-5) complex together with a target RNA oligonucleotide in the presence of ATP and magnesium ions at 37 °C. We subsequently heat-inactivated the complex, removed precipitated proteins by centrifugation and added a fraction of the deproteinized supernatant to an assay reaction mixture containing EiCsm6 and the fluorogenic RNA substrate. We observed robust activation of EiCsm6 by the supernatant, with 1% of the added supernatant having the same effect as 500 nM A6>P, suggesting that the EiCsm(1-5) complex produces a diffusible molecular product capable of activating EiCsm6 (Fig. 3a). Activator production was dependent on the presence of ATP, which could not be substituted by any of the three other nucleotide triphosphates, and required Mg^2+^ ions, as addition of EDTA was inhibitory (Fig. 3b). Moreover, the activator was only produced in the presence of a cognate target RNA but not when a non-cognate control RNA was used instead (Fig. 3b). Together, these results show that the EiCsm(1-5) complex generates a diffusible Csm6 activator by an ATP-, magnesium- and target RNA-dependent mechanism.

**Figure 3.**
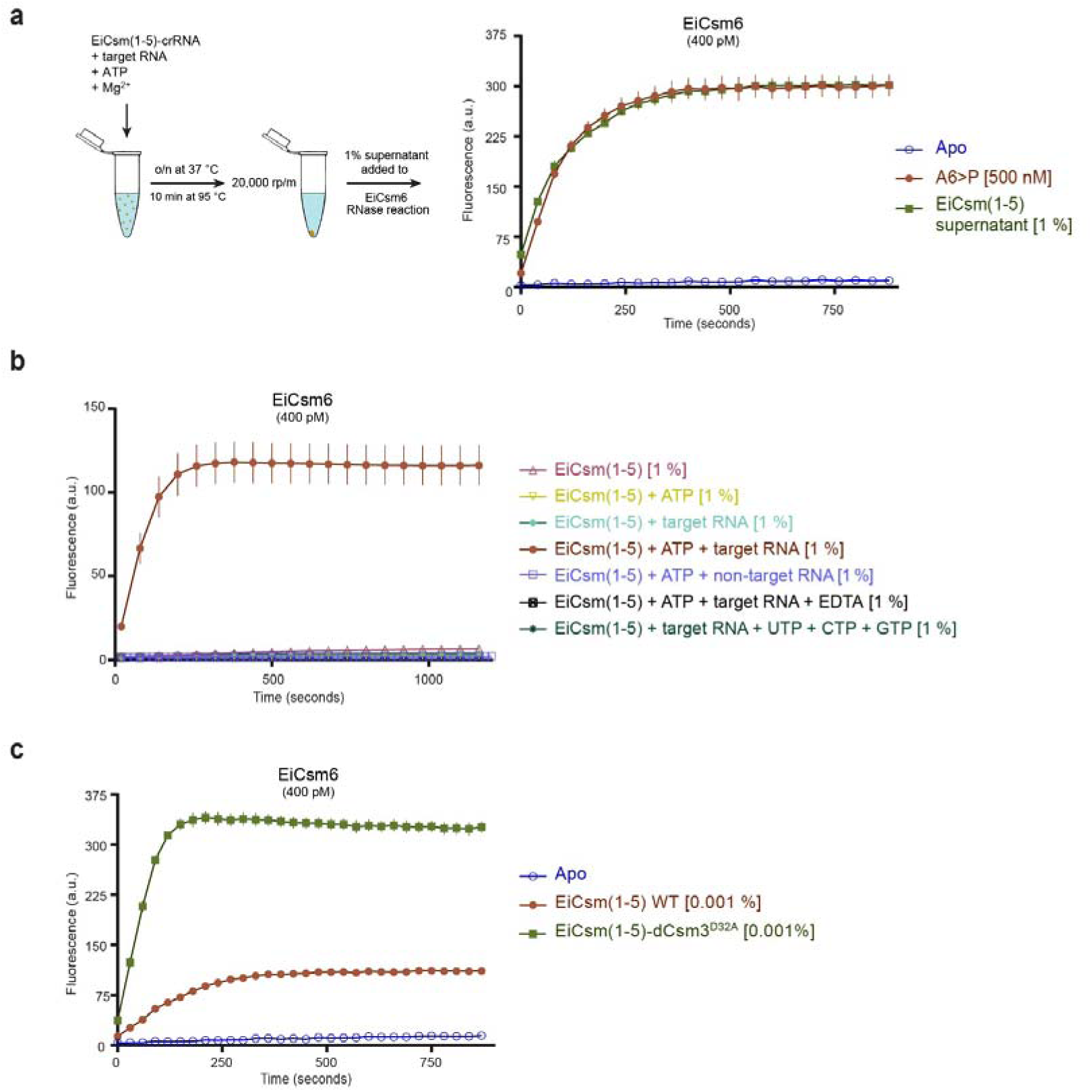
The Cas10 complex activates Csm6 via a diffusible second messenger. **a,** Left: Schematic representation of EiCsm(1-5) complex activation assay; right: RNase activity of EiCsm6 in the presence of synthetic A6>P or the product generated by the EiCsm(1-5) complex. **b**, Control assay to identify components necessary for activator production by the EiCsm(1-5) complex. **c**, EiCsm6 RNase activity in the presence of trace amounts of the product generated by the WT EiCsm(1-5) and EiCsm(1-5)-dCsm3^D32A^ complexes. All data points represent the mean of three replicates ± s.e.m.

Since crRNA-guided target RNA cleavage by the Csm(1-5) complex yields hexanucleotide products that could potentially act as Csm6 activators, we tested whether the ribonuclease activity of the EiCsm(1-5) complex was required to generate the EiCsm6 activator. Supernatant from a reaction containing EiCsm(1-5) complex in which the crRNA-guided RNase activity was abolished by the D32A mutation in the Csm3 subunits activated EiCsm6 even more strongly than the wild-type complex (Fig. 3c). These results indicate that activator generation is dependent on target RNA binding but activator is not generated by cleavage of target RNA. Moreover, the observed hyperactivity of the RNase-deficient EiCsm(1-5) complex indicates that production of the Csm6 activator is itself allosterically activated by target RNA binding by the type III interference complex.

## The Palm domain of Cas10 converts ATP into a cyclic oligoA product

Having ruled out the crRNA-guided target RNA cleavage activity as the source of the allosteric effector, we examined other enzymatic activities within the type III interference complex. The Cas10 (Csm1) subunit of the complex contains two putative catalytic domains. The histidine-aspartate (HD) domain of Cas10 possesses sequence non-specific DNase activity and mediates target RNA-dependent DNA degradation^9-12^. In turn, the Palm domain shares structural similarity with nucleotidyl cyclases and nucleotide polymerases and contains a putative, conserved Gly-Gly-Asp-Asp (GGDD) catalytic motif^27^. To test whether either of the domains was required for the production of the Csm6 activator, we generated EiCsm(1-5) complexes harbouring inactivating point mutations in the HD domain (HD>AN) or in the Palm domain motif (GGDD>GGAA) of Cas10. Whereas inactivation of the HD domain had no effect on the production of the Csm6 activator, mutations in the Palm domain completely abrogated activator formation (Fig. 4a), indicating that the Cas10 Palm domain generates the Csm6 activator.

**Figure 4.**
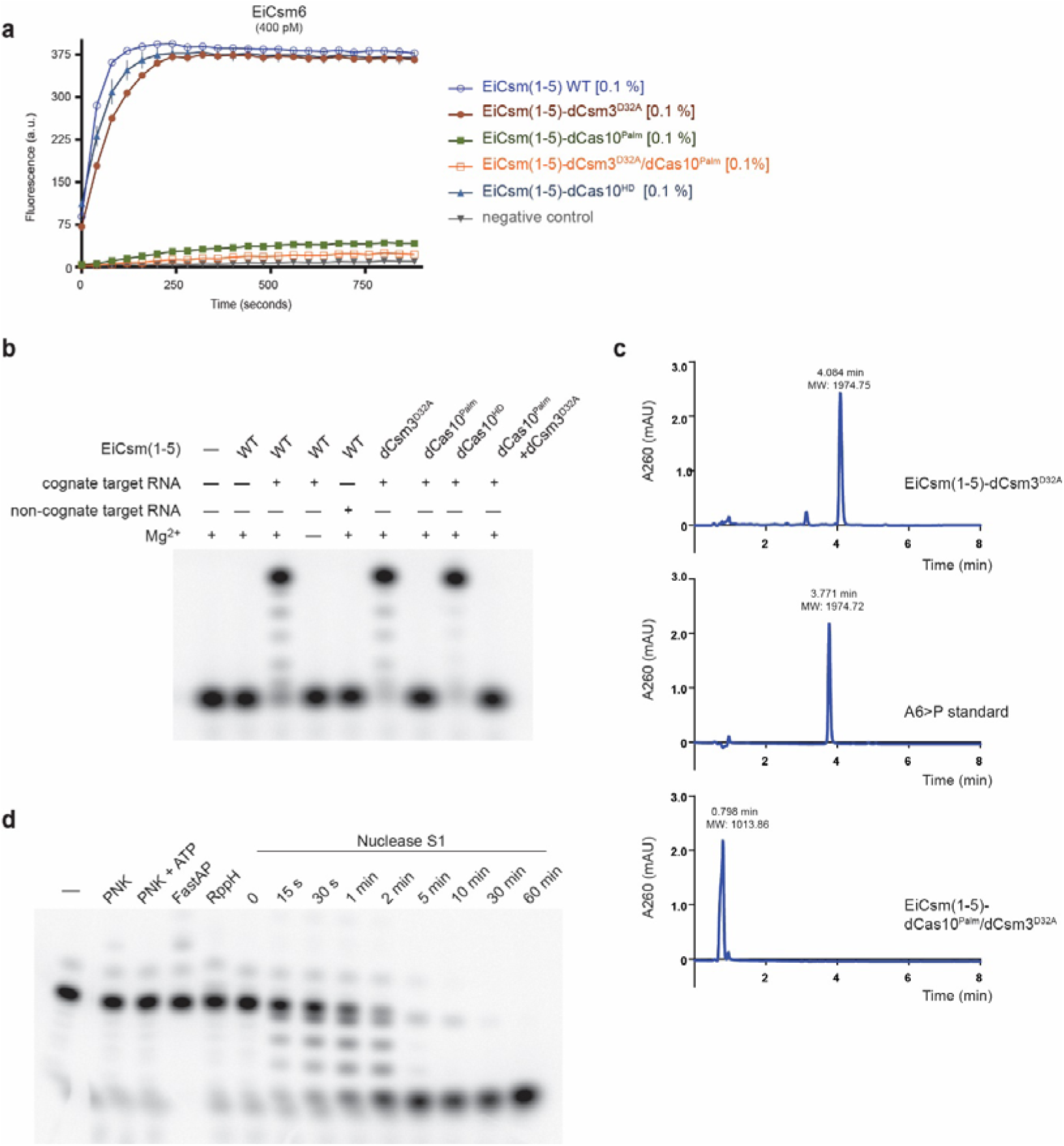
The Palm domain of Cas10 generates cyclic hexaadenylate *in vitro*. **a,** EiCsm6 RNase activity in the presence of products generated by WT and mutant EiCsm(1-5) complexes. **b,** ATP oligomerisation assay using [α-^32^P] ATP**. c,** Liquid chromatography-mass spectrometry (LC-MS) analysis of the products generated by the EiCsm(1-5)-dCsm3^D32A^ complex, the EiCsm(1-5)-dCas10^Palm^/dCsm3^D32A^ complex, and a synthetic A6>P standard. **d,** Treatment of the product generated by the EiCsm(1-5)-dCsm3^D32A^ complex by T4 polynucleotide kinase (T4 PNK), alkaline phosphatase (FastAP), pyrophosphatase RppH and S1 nuclease.

Previous structural studies of the Cas10 subunit (Cmr2) from the type III-B interference complex of *Pyrococcus furiosus* revealed that the putative Palm domain active site can accommodate two ATP molecules, while the GGDD motif coordinates divalent metal ions^28^. Based on this, we hypothesized that the Palm domain functions as an oligoA synthetase by directly catalysing 5’-3’ phosphodiester formation between ATP molecules. To test this possibility, we incubated the EiCsm(1-5) complex in the presence of target RNA, Mg^2+^ ions and [α-^32^P]-ATP, and visualized reaction products by denaturing polyacrylamide gel electrophoresis and phosphorimaging. In the presence of cognate target RNA, the EiCsm(1-5) complex converted ATP into a product displaying lower electrophoretic mobility. In contrast, product formation did not occur when a non-cognate RNA was supplied (Fig. 4b). Moreover, the activity was dependent on the presence of an intact GGDD motif in the Palm domain, whereas inactivating point mutations in the Cas10 HD domain or in Csm3 had no effect (Fig. 4b). Together, these results indicate that within the Csm(1-5) complex, the Cas10 Palm domain is allosterically activated by target RNA binding to generate a well-defined oligoA product by ATP polymerization.

Liquid chromatography-mass spectrometry (LC-MS) analysis revealed that the EiCsm(1-5) complex generates a single dominant molecular species with a molecular mass of 1974.75 Da, consistent with the product being either A6>P or a cyclic adenine hexanucleotide (Fig. 4c, Extended Data Fig. 6). However, the retention time of the product (4.08 min) was markedly different from that of a synthetic A6>P standard (3.77 min), indicating that the product is cyclic rather than linear (Fig. 4c). The product was refractory to treatments with polynucletide kinase (PNK), alkaline phosphatase and the pyrophosphatase RppH, indicating that it lacked 5’- or 3’-phosphate, 2’,3’-cyclic or 5’-triphosphate moieties. Digestion of the product with S1 nuclease yielded one terminal species, further confirming that the product is a cyclic, 3’-5’ linked hexanucleotide (Fig. 4d). These results indicate that the Palm domain of the Cas10 subunit of the *E. italicus* type III interference complex is a cyclic oligoadenylate synthetase that uses ATP to generate a cyclic hexaadenylate product.

Taken together, the results suggest that cyclic oligoadenylates are allosteric activators of Csm6 enzymes *in vivo* and that the activators are synthesized by the type III CRISPR machinery upon target RNA recognition. In light of our TtCsm6 data, these results also imply that other type III systems might generate cyclic tetranucleotides instead. Consistent with this possibility, a recent structure of the CARF domain protein Csx3 bound to a linear tetranucleotide shows that the ligand adopts a pseudocircular conformation^29^.

## *In vivo* activity of Csm6 is dependent on the oligoadenylate cyclase activity of the Csm complex

In the *S. epidermidis* type III-A CRISPR-Cas system, the RNase activities of Csm6 and Csm3 result in the degradation of phage transcripts, which is required for efficient anti-phage immunity when the target site is located in a gene transcript expressed late during infection^16^. To corroborate our biochemical studies, we tested whether the *in vivo* activity of Csm6 depends on allosteric activation by oligoA generated by Cas10. We first determined whether EiCsm6 can function together with the *S. epidermidis* type III system in the absence of its endogenous *csm6* gene. To this end, we engineered *Staphylococcus aureus* strains containing an RNase-deficient version of the *S. epidermidis* type III-A system that was programmed to target the *gp43* gene of phage ΦNM1γ6, contained an inactivating point mutation in the SeCsm3 protein and lacked SeCsm6. The strains co-expressed either wild-type EiCsm6 or the unresponsive EiCsm6 Q116A mutant (dEiCsm6^CARF^) from a plasmid. Cell cultures were infected with ϕNM1γ6 phage at low multiplicity of infection (MOI ~0.25) and cell survival was measured over time (Fig. 5a). Staphylococci expressing EiCsm6 were equally, if not more, immune as those expressing catalytically active SeCsm6, demonstrating that the SeCsm(1-5) complex can activate EiCsm6. When the dEiCsm6^CARF^ mutant was expressed instead, CRISPR-mediated immunity was lost and staphylococci succumbed to phage infection, as did control strains expressing the HEPN RNase active site mutants of SeCsm6 or EiCsm6 (Fig. 5a). These results suggest that oligoA sensing by the Csm6 CARF domain is required for Csm6 function *in vivo* and also affirm that Csm6 activation relies on a diffusible ligand rather than a direct physical interaction with the type III interference complex.

**Figure 5.**
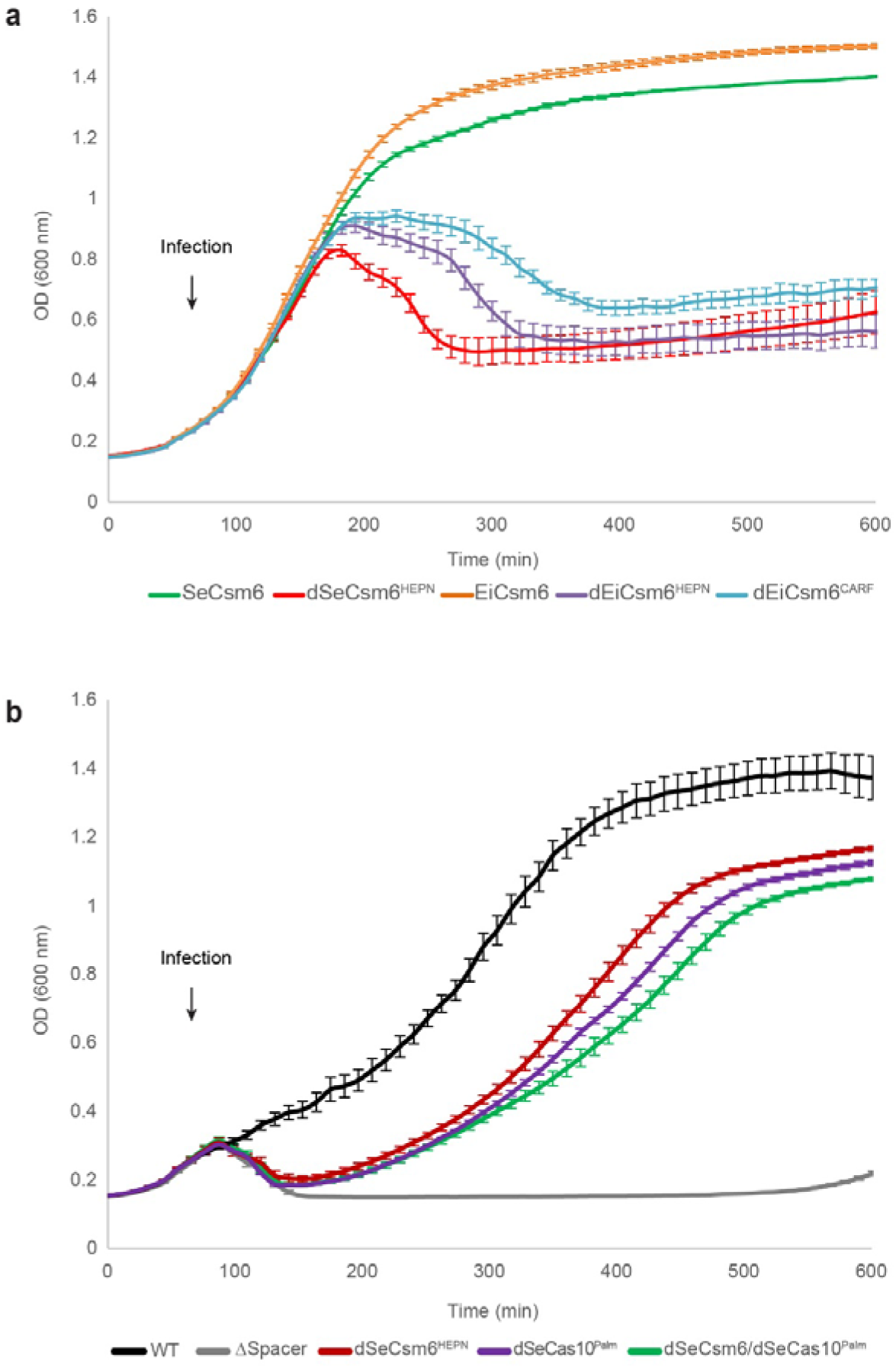
*In vivo* activity of Csm6 is dependent on cyclic oligoA and Cas10. **a,** Optical density of *S. aureus* cells containing RNase-deficient type III-A CRISPR-Cas system of *S. epidermidis* programmed with a *gp43* spacer. The system contains an inactivating point mutation in the *csm3* gene (dCsm3^D32A^) and a deletion of the *csm6* gene (ΔCsm6). A second plasmid expresses wild-type or mutated forms of *S. epidermidis* or *E. italicus* Csm6. Cells were infected at 60 minutes with bacteriophage ϕNM1γ6 at a multiplicity of infection (MOI) of 0.25. Each data points represents the mean of three replicates ± s.e.m. **b**, Growth curves of *S. aureus* strains harbouring the type III-A CRISPR system of *S. epidermidis* with a *gp43* spacer and indicated *cas* gene mutations. Infection with ϕNM1γ6 is initiated at 60 minutes with an MOI of 30. Each data points represents the mean of three replicates ± s.e.m.

We then proceeded to test whether the function of the endogenous Csm6 protein is dependent on the enzymatic activity of the Palm domain of Cas10. To this end, we infected staphylococci containing the *S. epidermidis* system programmed to target the late-expressed gene *gp43* with phage ϕNM1γ6. Whereas the activity of either Csm3 or Csm6 is sufficient to support immunity at low multiplicity of infection (MOI ~1)^16^, at high MOI (~30) mutation of the Csm6 HEPN RNase domain (dCsm6^HEPN^) resulted in a significant reduction of cell survival, even in the presence of active Csm3 (Fig. 5b). Mutation of the GGDD motif in the Cas10 subunit (dSeCas10^Palm^) resulted in similar reduction in cell survival as that observed for the dSeCsm6^HEPN^ mutant (Fig. 5b). It was previously reported that this mutation in *S. epidermidis* Cas10 prevents DNA cleavage *in vitro*^6^, and therefore the defect in immunity could be attributed to a functional role of the Palm domain in Csm6 activation, DNA cleavage or both. To distinguish between these possibilities we additionally tested a dSeCsm6^HEPN^/dSeCas10^Palm^ double mutant strain, reasoning that the combined effect of the mutations would not be additive if immunity is a consequence of Cas10-dependent Csm6 activation. Indeed, the double mutant displayed a cell survival profile similar to each of the individual mutants, indicating that Cas10 Palm domain and Csm6 RNase activity are genetically linked. Altogether, these results demonstrate that the *in vivo* activity of Csm6 is dependent on the cyclic oligoadenylate synthetase activity of the type III CRISPR interference complex.

## Discussion

Previous studies demonstrated that Csm6 family ribonucleases provide an additional interference mechanism in type III CRISPR-Cas systems by targeting invader transcripts^16^. Here we show that Csm6 proteins are allosterically activated by cyclic oligoA effectors through binding to the CARF domain and that the effector is synthesized by the Palm domain of the Cas10 subunit of the type III interference complex in response to target RNA recognition. Our findings demonstrate that type III systems use second messengers to signal target RNA sensing to CARF domain ribonucleases. Although bacteria utilize numerous nucleotide metabolites such as c-di-AMP, c-di-GMP or ppGpp for signalling and stress response pathways^30^, cyclic oligoA metabolites have not been reported to function as second messengers in prokaryotes before. However, the mechanism brings up striking parallels with the vertebrate innate immune system, where detection of viral double-stranded RNAs by oligoadenylate synthetase enzymes triggers production of linear 5’-2’ linked oligoadenylates that allosterically activate ribonuclease L^31^.

We conclude that the type III interference complex is not only a crRNA-guided RNase and DNase, but also a cyclic oligoadenylate synthetase (Fig. 6). The likely mechanism of cyclic oligoA synthesis by Cas10 would involve linear oligomerization, followed by final cyclization of the linear intermediate through its 5’-triphosphate group (Extended Data Fig. 7). As the cyclic oligoadenylate synthetase activity is stimulated by target RNA binding, this would provide a fail-safe interference mechanism in case the intrinsic DNase and RNase activities of the type III interference complex are insufficient to clear the invader DNA and/or its transcripts, such as when the target gene is expressed late in the phage lytic cycle or contains mismatches to the crRNA guide^16^. Notably, CARF domain ribonucleases of the Csm6 and Csx1 families are only associated with type III CRISPR-Cas systems containing a Cas10 subunit with a catalytically competent Palm domain. This hints that the second messenger-mediated activation mechanism is nearly universally conserved in type III-A and III-B CRISPR-Cas systems. Moreover, the existence of other CRISPR-associated CARF domain proteins, such as Csa3^13,32^, that appear to be transcription factors rather than nucleases raises the possibility that CRISPR-Cas systems also use cyclic oligoA signalling for transcriptional regulation, either within their own loci or to induce other genome defence mechanisms or stress response pathways upon invader detection.

**Figure 6.**
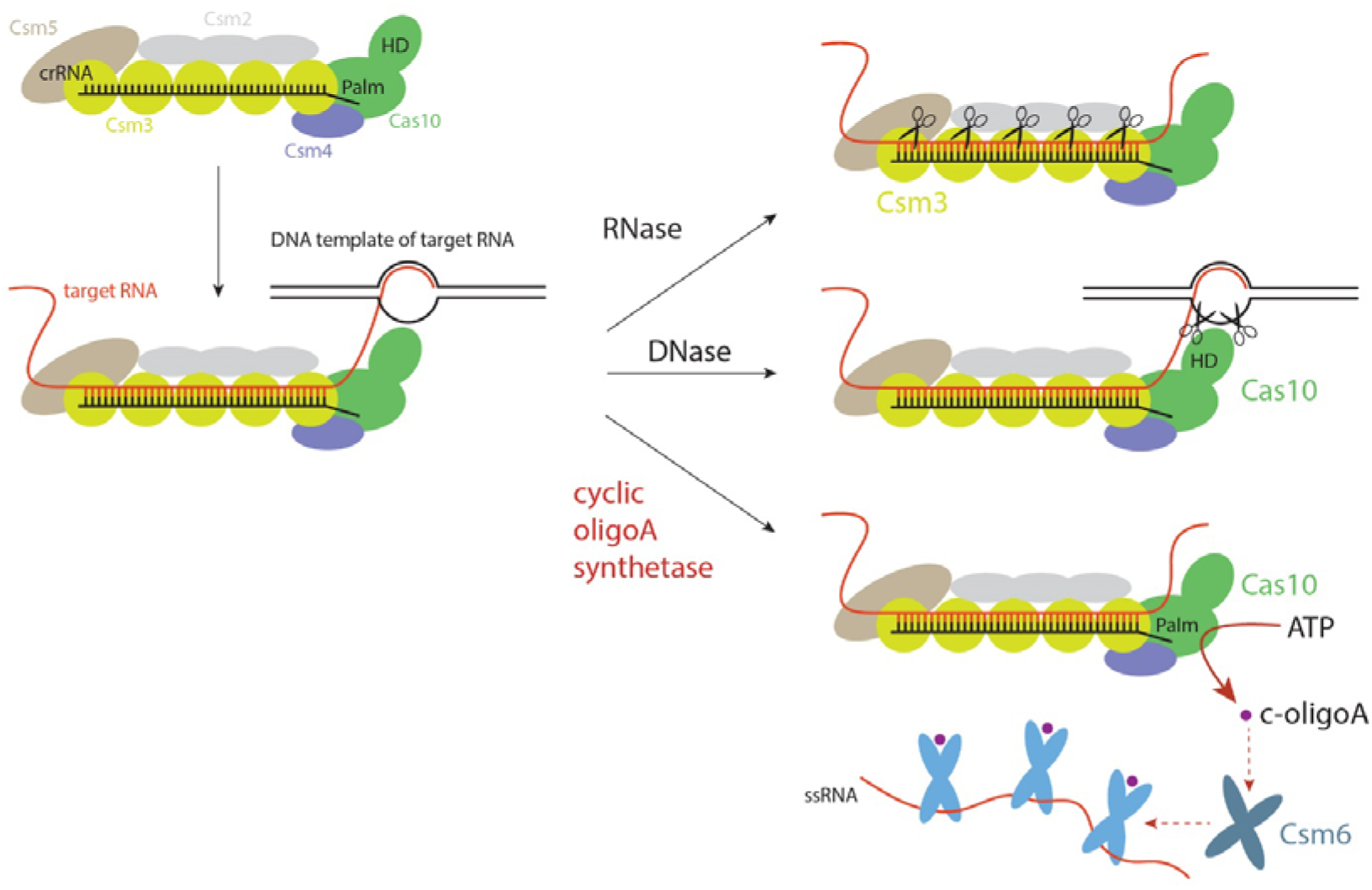
Proposed model for the molecular mechanism of type III CRISPR-Cas systems. The type III CRISPR interference complex has three enzymatic activities: (i) a crRNA-guided endoribonuclease activity against target RNA harboured by the Csm3 subunits, (ii) target RNA-stimulated DNase activity harboured by the HD domain of Cas10 and (iii) target RNA-stimulated cyclic oligoadenylate synthetase activity harboured by the Palm domain of Cas10. The cyclic oligoA product of Cas10 allosterically activates Csm6 RNase.

## AUTHOR CONTRIBUTIONS

O.N, C.G-D. and M.J. conceived the study. O.N., C.G.-D., J.T.R., L.M. and M.J. designed experiments. O.N. expressed and purified recombinant Csm6 proteins, carried out oligoA activation and ATPase assays and performed enzymatic probing of the cyclic oligoA product. C.G.-D. expressed and purified recombinant EiCsm(1-5) complexes, performed oligoA activation assays and assisted with LC-MS analysis. J.T.R. performed phage infection assays under supervision of L.M. C.B. synthesized 2’,3’-cyclic phosphate-terminated nucleotides and carried out LC-MS analysis under supervision of J.H. F.S. synthesized 2’,3’-cyclic phosphate-terminated A4 nucleotide and advised on nucleotide chemistry. L.B. performed additional LC-MS analyses of Csm6 effectors. O.N., C.G.-D. and M.J. wrote the manuscript, with input from the remaining authors.

## ACKNOWLEDGEMENTS

We thank members of the Jinek and Marraffini laboratories for helpful discussions and critical comments on the manuscript. We thank Daan Swarts for technical assistance and sharing reagents. We thank Undine Manzau for technical assistance during A4>P synthesis and purification. This study was supported by a Swiss National Science Foundation project grant to M.J. (SNSF 31003A_149393) and by funding from the Swiss National Competence Center for Research (NCCR) “RNA & Disease” (to M.J. and J.H.). M.J. is International Research Scholar of the Howard Hughes Medical Institute and Vallee Scholar of the Bert L & N Kuggie Vallee Foundation. C.G.-D. was supported by a Long-Term Fellowship from the European Molecular Biology Organization (EMBO). J.T.R. was supported by a Boehringer Ingelheim Fonds PhD fellowship. L.A.M. is supported by the Rita Allen Scholars Program, a Burroughs Wellcome Fund PATH award, an NIH Director’s New Innovator Award (1DP2AI104556-01) and a HHMI-Simons Faculty Scholar Award.

## CONFLICT OF INTEREST

The authors declare no conflicts of interest.

## METHODS

### Plasmid DNA constructs

DNA sequences encoding for TtCsm6 (*Thermus thermophilus* DSM-579), EiCsm6 (*Enterococcus italicus* DSM-15952) and MtCsm6 (*Methanothermobacter thermoautotrophicus* DSM-1053) were amplified from their respective genomic DNAs by a standard PCR protocol and were subsequently cloned into a 2HR-T vector (Addgene #29718) using ligation-independent cloning, resulting in constructs carrying an N-terminal hexahistidine tag followed by a StrepII tag, a Tobacco Etch Virus (TEV) protease cleavage site, and the Csm6 polypeptide sequence. To obtain a plasmid vector for heterologous expression a functional EiCsm(1-5) interference complex from the *E. italicus* DSM-15952 type III-A CRISPR locus, a DNA fragment spanning from the *cas10* gene to the end of the second spacer (GenBank GL622241.1; complement 1999065-1991480) was amplified from *E. italicus* genomic DNA using primers oCGD208 and oCGD209 and cloned into a p7XNH3 vector using FX cloning^36^, resulting in the pCGD253 plasmid. In this plasmid, Cas10 (Csm1) was N-terminally fused downstream of a deca-histidine tag and a human rhinovirus 3C protease cleavage site. Point mutations in all constructs were inserted by inverse PCR using the primers and templates listed in Supplementary Table 1, and resulting plasmids were verified by DNA sequencing. Synthetic DNA and RNA oligonucleotides were obtained from Sigma-Aldrich or IDT and are listed in Supplementary Table 2.

The plasmids containing the wild-type *S. epidermidis* RP62a CRISPR system with an anti-gp43 spacer and wild type SeCsm6 (pWJ191) or inactive dCsm6^HEPN^ (R364A, H369A) mutant (pWJ241), as well as the no-spacer control plasmid (pGG-BsaI-R) were previously described^16^. To construct the Cas10 Palm GGAA (D586A, D587A) mutation (dCas10^Palm^), the wild-type plasmid was amplified by PCR with primer pairs W852/W1169 and W614/W1170, and the resulting fragments were ligated by Gibson assembly. The plasmid carrying the dCsm6^HEPN^ (R364A, H369A) and dCas10^Palm^ double mutation (pJTR147) was generated by amplifying PCR products from pWJ241 with primers W614/PS93 and from pWJ299 with primers W852/PS94, and them by Gibson assembly. pWJ235 was generated by amplifying PCR products from the plasmid pWJ192 with primer pairs PS465/W1022 and W494/PS466, and joining them via Gibson assembly. pWJ156, containing SeCsm6, was constructed by a one-piece Gibson reaction using the PCR product from the amplification of pAS9 with primers W795 and W796. The plasmid carrying dCsm6^HEPN^ was generated by amplifying pWJ156 with NP38+PS247 and NP39+PS248, and joining the products by Gibson ligation. The plasmids containing wild type EiCsm6 (pJTR179), dEiCsm6^HEPN^ (pJTR180), and dEiCsm6^CARF^ (pJTR181) were generated by amplifying DNAs from plasmids pOP214, pOP215, or pOP218 respectively with primers JTR504/JTR505. Each resulting fragment was combined by Gibson ligation with the PCR product amplified from pWJ156 with primers JTR502/JTR503.

### Protein expression and purification

For expression of TtCsm6, MtCsm6, EiCsm6, and EiCsm(1-5) complex, the corresponding plasmid was transformed into *E. coli* BL21 Rosetta2 (DE3) cells. Cells were grown in LB supplemented with appropriate antibiotics until they reached OD_600nm_ ~0.6 and expression was induced by addition of 0.2 mM IPTG (isopropyl-β-D-thiogalactopyranoside). Proteins were expressed at 18 °C for 16 h. For TtCsm6, MtCsm6 and EiCsm6 the cells were harvested and resuspended in lysis buffer containing 20 mM HEPES pH 8.0, 500 mM KCl, and 5 mM imidazole pH 8.0. The cell suspension was lysed by ultrasonication and lysate was cleared by centrifugation at 20,000 g for 40 min. Cleared lysate was applied to a 5 ml Ni-NTA cartridge (Qiagen), the column was washed with 10 column volumes of lysis buffer and bound proteins were eluted with 5 column volumes of the same buffer supplemented with 250 mM imidazole pH 8.0. Eluted proteins were dialysed overnight against 20 mM HEPES pH 7.5 and 500 mM KCl in the presence of TEV (tobacco etch virus) protease. Dialysed proteins were passed through a 5 ml Ni-NTA cartridge. The flow-through fraction was concentrated and further purified by size-exclusion chromatography using an S200 (16/600) column (GE Healthcare) in 20 mM HEPES pH 7.5 and 500 mM KCl, yielding pure, monodisperse proteins. Purified proteins were concentrated to 5-75 mg ml^-1^ using 30,000 MWCO centrifugal filters (Merck Millipore) and flash-frozen in liquid nitrogen.

For EiCsm(1-5) complex, cells were harvested and resuspended in lysis buffer containing 20 mM HEPES pH 8.0, 400 mM KCl, and 5 mM imidazole supplemented with EDTA-free protease inhibitor (Roche) and lysed using an HPL6 cell homogenizer. Lysate was clarified by centrifugation at 20,000 g for 40 min. The lysate was applied to a 5 ml Ni-NTA cartridge (Qiagen), the column was washed in two steps with lysis buffer supplemented 10 and 50 mM imidazole. The complex was eluted with elution buffer containing 20 mM HEPES pH 7.5, 400 mM KCl, and 250 mM Imidazole. The complex was concentrated and further directly purified by size-exclusion chromatography using two Superose-6 (10/300) columns connected in tandem, or a Sephacryl S-300 (26/600) column (GE Healthcare), in buffer containing 20 mM HEPES pH 7.5, 300 mM KCl, and 5 mM MgCl_2_. Appropriate fractions were pooled and concentrated using 100,000 MWCO centrifugal filters (Merck Millipore), and samples were analysed by SDS-PAGE using a 4-15% gradient polyacrylamide gel (Bio-Rad). All mutant complexes were purified using the same protocol as for the wild type complex. The concentration of EiCsm(1-5) complex was calculated according to a formula that takes nucleic acid contamination (i.e. crRNA) into consideration.

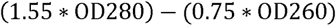

### Synthetic RNA oligonucleotides

A synthetic target oligonucleotide for the EiCsm(1-5) was obtained from IDT with the sequence 5’- GGGUAAGGAUAGUGAAAUAGAUGCAGUGGAAGCAUACCAAGAGCA-3’. A non-cognate control RNA contained the sequence 5’- GGTCTATTGCCCTCTATATCGGGCTGTTCTCCAAACCGGTCTGGGTACCTTCGTACGGACGACCTTC-3’.

Linear 3’-hydroxylated oligoA nucleotides were obtained from Dharmacon. Linear tetraadenylate carrying a 3’-phosphate group (ApApApAp) was obtained from Biospring AG, and subsequently converted to A4>P as follows. 2.5 ml 2-(N-morpholino)ethanesulfonic acid buffer (MES, 20 mM, pH 6) and 1-Ethyl-3-(3-dimethylaminopropyl)carbodiimide (EDC, 5.9 mg, 38 μmol, 3.1 eq.) were added to ApApApAp (12.3 μmol in 486 μl water) and the resulting solution was stirred at room temperature. Further portions of 1 ml MES-buffer and 1.91 mg (1 eq.) EDC were added to the reaction mixture after 24 hours and 48 hours. HPLC analysis after 55 hours indicated ~90 % product formation. The reaction mixture was diluted with 10 ml water and was extracted with chloroform (4 x 40 ml). The combined organic phases were back-extracted with 10 ml water and the combined product-containing aqueous phase was filtered (regenerated cellulose membrane, 5 μm) to remove particulate components. The product solution was diluted with water to ~90 ml and applied to a Q Sepharose Fast Flow anion exchange column (40–165 μm; 100 x 10 mm), previously regenerated with 2 M sodium chloride and washed with water. The column was washed with water, followed by a gradient of 0-2 M NaCl. The title compound eluted with ~0.2–0.3 M NaCl. Product-containing fractions were carefully concentrated to a final volume of approximately 10 ml. Desalting of A4>P was accomplished by semi-preparative reverse-phase HPLC. The product solution was applied to a YMC*GEL ODS-A 12 nm column (10 μm; 250 x 16 mm) pre-equilibrated with 100 % water. The column was washed with water to remove excess NaCl. 50 % methanol in water was used to elute residual title compound. Product-containing fractions of appropriate purity were pooled, concentrated with a rotary evaporator and subsequently lyophilised in high vacuum for 16 hours to yield 5.8 μmol A4>P, sodium salt (theoretical yield: 47.2 %; purity: 99.96 % by HPLC).

All other linear oligoA nucleotides containing 2’,3’-cyclic phosphate groups (A5>P, A6>P, A7>P, A8>P) were synthesized as follows. Phosphoramidites were obtained from Thermo Fisher Scientific (Waltham, MA, USA). 2’,3’-cyclic phosphate oligonucleotides were synthesized on a MM12 synthesizer from Bio Automation (BioAutomation Corp., Irving, TX, USA) using 5 mg 5’-DMT-2’, 3’-Cyclic Phosphate Adenosine (N-Bz) 1000 Å CPG (ChemGenes, Wilmington, MA, USA) and 2’-TBDMS Adenosine (N-Bz) CED phosphoramidite with a coupling time of 2 x 90 s. Phosphoramidite was prepared as a 0.08 M solution in dry acetonitrile (ACN), the activator 5-(Benzylthio)-1H-tetrazole (Biosolve BV, Valkenswaard, Netherlands) was prepared as a 0.24 M solution in dry ACN. Oxidizer was prepared as a 0.02 M I_2_ solution in THF/pyridine/H_2_O (70:20:10, w/v/v). Capping reagent A was THF/lutidine/acetic anhydride (8:1:1) and capping reagent B was 16% Nmethylimidazole/THF. 3% dichloroacetic acid in dichloromethane was used as deblock solution. 2’,3’-cyclic phosphate oligonucleotides were synthesized DMT-off. After completion of the automated synthesis, oligonucleotides were cleaved from the solid support and the protecting groups on the exocyclic amino groups (N6-benzoyl) and the backbone (2-cyanoethyl) were removed using a 1:1 mixture of 40% aqueous methylamine and 25% aqueous ammonia for 1 h at 65 °C. 2’-O-tert-butyldimethylsilyl groups (TBDMS) were removed and simultaneously the 2’,3’-cyclic phosphate terminus was formed using a fresh mixture of N-methyl-2-pyrrolidone (60 μl), triethylamine (30 μl) and triethylamine trihydrofluoride (40 μl) at 40°C for 6 h. Desilylation was quenched with trimethylethoxysilane (200 μl), and diisopropyl ether (200 μl) was subsequently added to precipitate the oligonucleotide. The precipitate was dissolved in H_2_O and purified on an Agilent 1200 series preparative HPLC fitted with a Waters XBridge Oligonucleotide BEH C18 column (10 × 50 mm, 2.5 μm) at 65 °C using a gradient of 5-20% buffer B over 8 min and flow rate of 5 ml min^-1^. Buffer A was 0.1 M triethylammonium acetate pH 8.0; buffer B was methanol. Fractions were pooled, dried under vacuum and dissolved in H_2_O.

### Gel-based Csm6 ribonuclease activity assay

Gel-based cleavage assays were performed as described previously^17^. Briefly, a 24-nucleotide synthetic RNA (250 nM), 5’-end labelled with Cy5, was incubated with Csm6 in the presence of indicated concentrations of oligonucleotide activators at 37 ^o^C. At indicated time points, the reaction was quenched by addition of an equal volume of quencher solution (95 % formamide, 5 % glycerol and 0.005 % bromophenol blue) and products were resolved on a 16 % denaturing (7M urea) polyacrylamide gel and visualised using a fluorescence gel scanner (Typhoon FLA 9500, GE Healthcare). Protein concentrations are indicated in the respective figure panels.

### Fluorescence-based Csm6 ribonuclease activation assay

To obtain kinetics of Csm6 ribonuclease activity, 2 μl of 2 μM RNase Alert substrate (IDT) and 1 μl of of synthetic oligoA or supernatant from a EiCsm(1-5) cyclic oligoadenylate synthetase reaction were added to a well in a 96-well plate. The reaction was started by addition of 97 μl Csm6 protein in 20 mM HEPES pH 7.5 and 50 mM KCl. Fluorescence signal (excitation at 490 nm, emission at 520 nm) was measured over time in TECAN Infinite M1000 or PHERAstar FSX (BMG Labtech) multimodal plate readers. Final protein and ligand concentrations used in the assay reactions are indicated in corresponding figure panels.

### Calculation of EC_50_

Cleavage kinetics for reactions containing EiCsm6 supplemented with indicated concentrations of A6>P were recorded and initial velocities were calculated over the first 10 s of the reaction. Initial velocities were then plotted against ligand concentrations in GraphPad and fitted by using a nonlinear regression analysis using the log(dose)-versus-response relationship, assuming a Hill coefficient of 1.

### In vitro cyclic oligoadenylate synthetase assays using EiCsm(1-5)

To utilise the EiCsm(1-5) complex and mutants for *in vitro* synthesis of cyclic oligoadenylates, 0.15 mg ml^-1^ EiCsm(1-5) complex was mixed with 10 μM target RNA, 1 mM MgCl_2_ and 0.5 mM ATP in a buffer containing 20 mM HEPES pH 7.5 and 50 mM KCl in a total volume of 30 μl. The reaction was incubated at 37 °C for ~16 h and the proteins were subsequently denatured at 95 °C for 10 min. Denatured protein was pelleted by centrifugation and the deproteinized supernatant was transferred to a fresh tube. The supernatant was diluted as indicated and added to the Csm6 ribonuclease activity assay as described above.

To directly visualize the synthesis of cyclic oligodenylates, the same reaction was supplemented with 1 μCi of [α-^32^P]-ATP (Hartmann Analytic GmbH) and the reaction time was shortened to 30 min. The reaction was stopped by heat-inactivation at 95 °C for 10 min. The products were mixed with equal volumes of quencher solution (95 % formamide, 5 % glycerol and 0.005 % bromophenol blue) and subsequently resolved on a 20 % denaturing (7M urea) polyacrylamide gel run at 60 W for ~5 h. The gel was dried and products were detected by phosphorimaging using a Typhoon FLA 9500 imager (GE).

### Nuclease S1 digest of Ei(Csm1-5) product

The radiolabelled product from the EiCsm(1-5) assay was digested with Nuclease S1 (Thermo Fisher Scientific). Cleavage reactions contained 4 μl 5x Nuclease S1 buffer, 15 μl of [α-^32^P] ATP-labelled oligoadenylates and 1 μl of Nuclease S1 diluted 1:50 in its commercial protein storage buffer (final concentration 0.1 U μl^-1^) in a total volume of 20 μl. The reaction was incubated at 37 °C and 2 μl aliquots were taken at indicated time points. Reactions were quenched by addition into 10 volumes quencher solution and subsequently resolved and imaged as described above.

### Enzymatic probing of Ei(Csm1-5) product

Aliquots of the radiolabelled product from the EiCsm(1-5) assay were treated with PNK (NEB) in the absence and presence of 1 mM ATP, with Fast Alkaline Phosphatase (Thermo Fisher Scientific) and *E.coli* RNA 5’ Pyrophosphohydrolase (RppH, purified in-house). All reactions were performed in reaction buffers optimised for the respective enzymes as recommended by the supplier. All reactions were performed in total volumes of 10 μl containing 1 μl enzyme, 1.5 μl [α-^32^P] ATP-labelled oligoadenylates, 1 μl 10x buffer and 6.5 μl H_2_O. Reactions were incubated at 37 °C for 90 min and products were resolved and imaged as above.

### LC-MS analysis

To analyse the products generated by EiCsm(1-5) complex, the EiCsm(1-5)-dCsm3^D32A^ complex was used to prevent cleavage of the target RNA by the Csm3 subunits and ensure persistent activation of the complex. A reaction containing 0.15 mg ml^-1^ of EiCsm(1-5) complex (500 nM), 10 μM target RNA, 1 mM MgCl_2_ and 0.5 mM ATP in buffer containing 20 mM HEPES pH 7.5 and 50 mM KCl was incubated for 30 minutes at 37 °C. To quench the reaction, the reactions were heated at 95 °C for 10 min, the denatured protein was removed by centrifugation and 1 μl of the deproteinized supernatant, diluted in 30 μl of H_2_O, was subjected to LC-MS. Analysis of purified oligonucleotides and enzymatic assays was conducted on an Agilent 1200/6130 LC-MS system fitted with a Waters Acquity UPLC OST C18 column (2.1×50 mm, 1.7 μm) at 65 °C, with a gradient of 5-20% buffer B in 10 min with a flowrate of 0.3 ml min^−1^. Buffer A was aqueous hexafluoroisopropanol (0.4 M) supplemented with triethylamine (15 mM). Buffer B was methanol. The supernatants obtained from the EiCsm(1-5)-dCsm3^D32A^ complex or the EiCsm(1-5)-dCsm3^D32A^/dCas10^Palm^ complex were analysed using the same protocol.

### RNA cleavage assay by EiCsm(1-5) complexes

The reaction was initiated by adding 250 nM target RNA substrate (3’-labelled with Alexa 352) to 250 nM EiCsm(1-5) complex (wild type or inactive mutants) in a buffer containing 20 mM HEPES pH 7.5 and 50 mM KCl. After 1 minute at 37 °C, the reaction was stopped by addition of equal volume of quencher solution (90% formamide, 5% glycerol, 25 mM EDTA) and heated at 95 °C for 5 min. 10% of the final reaction was resolved on a 16% denaturing (7M urea) polyacrylamide gel. The products were visualized using a Typhoon FLA 9500 fluorescence gel scanner.

### Phage infection assays

*S. aureus* RN4220 ^37^ was grown in tryptic soy broth (TSB) medium at 37°C, and supplied with 10 μg ml^-1^ chloramphenicol or 10 μg ml^-1^ erythromycin for plasmid maintenance. During bacteriophage infection, 5 mM CaCl_2_ was added to the media. For both EiCsm6 complementation and Csm6-Cas10 phenocopy experiments, overnight cultures initiated from single colonies were diluted 1:20 in TSB broth. After growth for 1 h, cell densities were normalised and cultures were added to a 96-microwell plate (Cellstar, 655180) in the presence of appropriate antibiotics and 5 mM CaCl_2_. The plate was shaken in a microplate reader (TECAN Infinite 200 PRO) at 37°C, with optical measurements taken at 600 nm every 10 minutes. Bacteriophage ϕNM1γ6^7^ was added at an MOI of approximately 0.25 or 30 after 60 min of shaking. For the complementation experiment, cells containing plasmid pWJ235 (gp43 spacer, dCsm3, ΔCsm6) in combination with one of the following plasmids; pWJ156, pJTR67, pJTR179, pJTR180, or pJTR181, were inoculated from single colonies and grown in TSB supplemented with 10 μg ml^-1^ chloramphenicol and 10 μg ml^-1^ erythromycin. For the phenocopy experiment, overnight cultures of *S. aureus* RN4220 containing pWJ191, pWJ241, pWJ299, pJTR147, or pGG-BsaI-R were inoculated from single colonies and grown in in TSB supplemented with 10 μg ml^-1^ chloramphenicol.

## Extended Data Figures

**Extended Data Figure 1.**
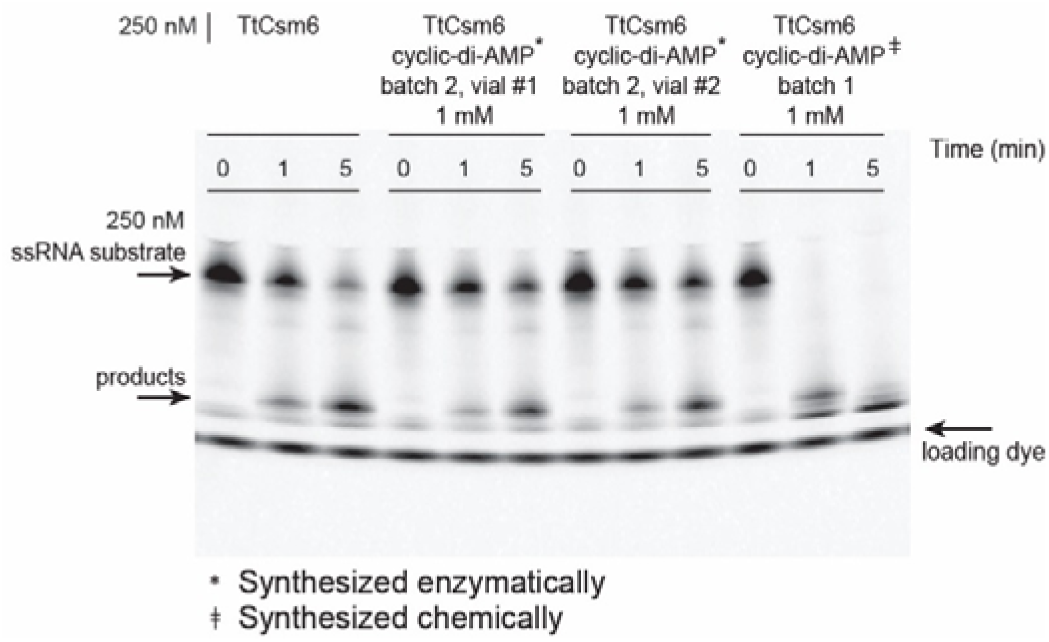
Activation of TtCsm6 by c-di-AMP contaminant. TtCsm6 RNase activity assay using a Cy5-labelled ssRNA substrate, in the presence of chemically or enzymatically synthesized c-di-AMP. TtCsm6 was incubated with 1 mM c-di-AMP from different batches and 250 nM of 5’-end Cy5 labelled ssRNA substrate at 37 °C and samples were taken at indicated time points. Reactions were quenched by addition of 1 volume of formamide. The reaction products were resolved by electrophoresis on a 16% denaturing (7M urea) polyacrylamide gel and visualised using a fluorescence gel scanner.

**Extended Data Figure 2.**
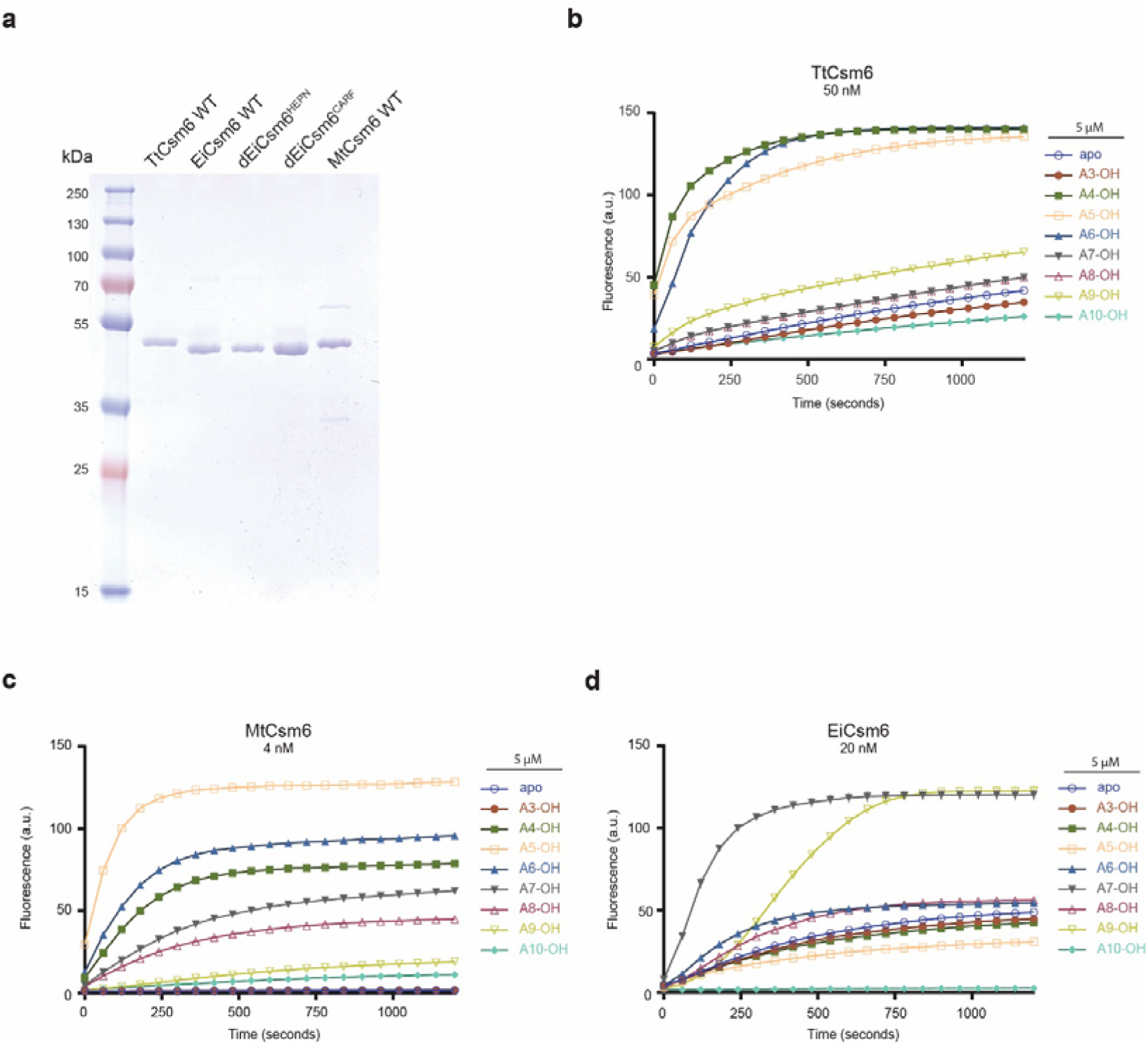
Activation of Csm6 ribonucleases by oligoadenylates. a, SDSPAGE analysis of purified WT and mutant Csm6 proteins. **b,c,d**, TtCsm6, MtCsm6, and EiCsm6 ribonuclease activity assays in the presence of 3’-hydroxylated oligoadenylates ranging from A3 to A10. All data points represent the mean of three replicates ± s.e.m.

**Extended Data Figure 3.**
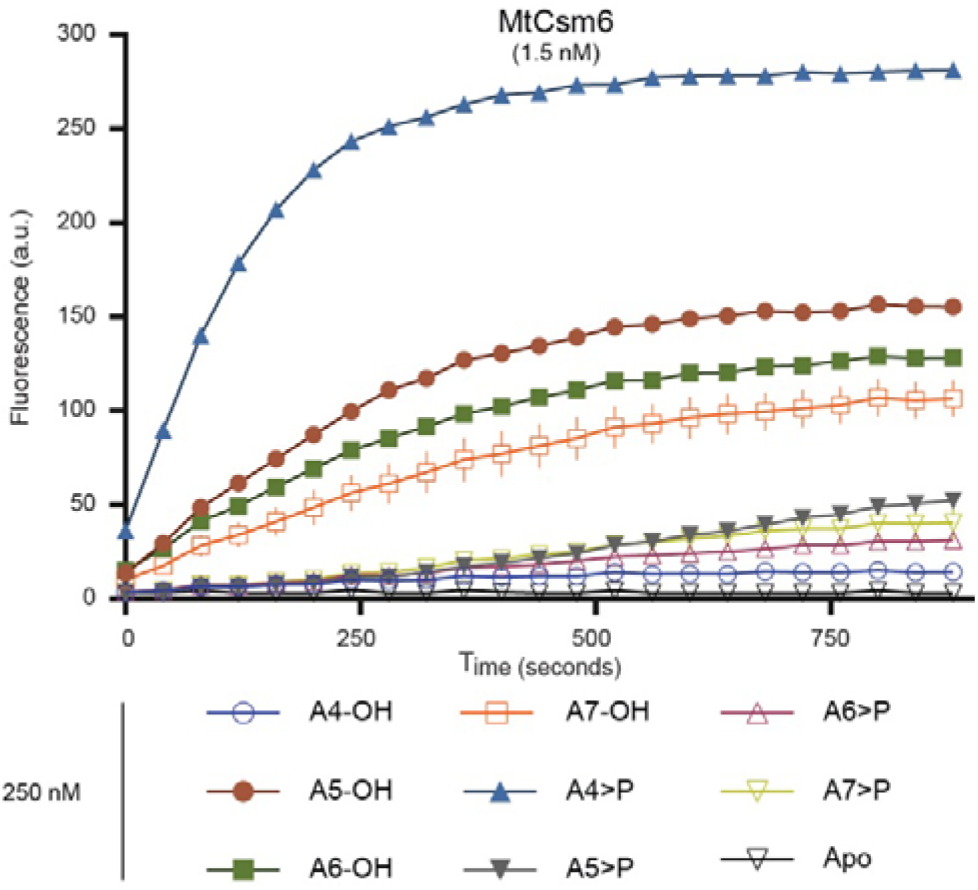
Activation of MtCsm6 by oligoadenylates with 2’,3’-cyclic phosphate. Ribonuclease activity assay in the presence of oligoadenylates carrying 3’- hydroxyl or 2’,3’-cyclic phosphate groups. All data points represent the mean of three replicates ± s.e.m.

**Extended Data Figure 4.**
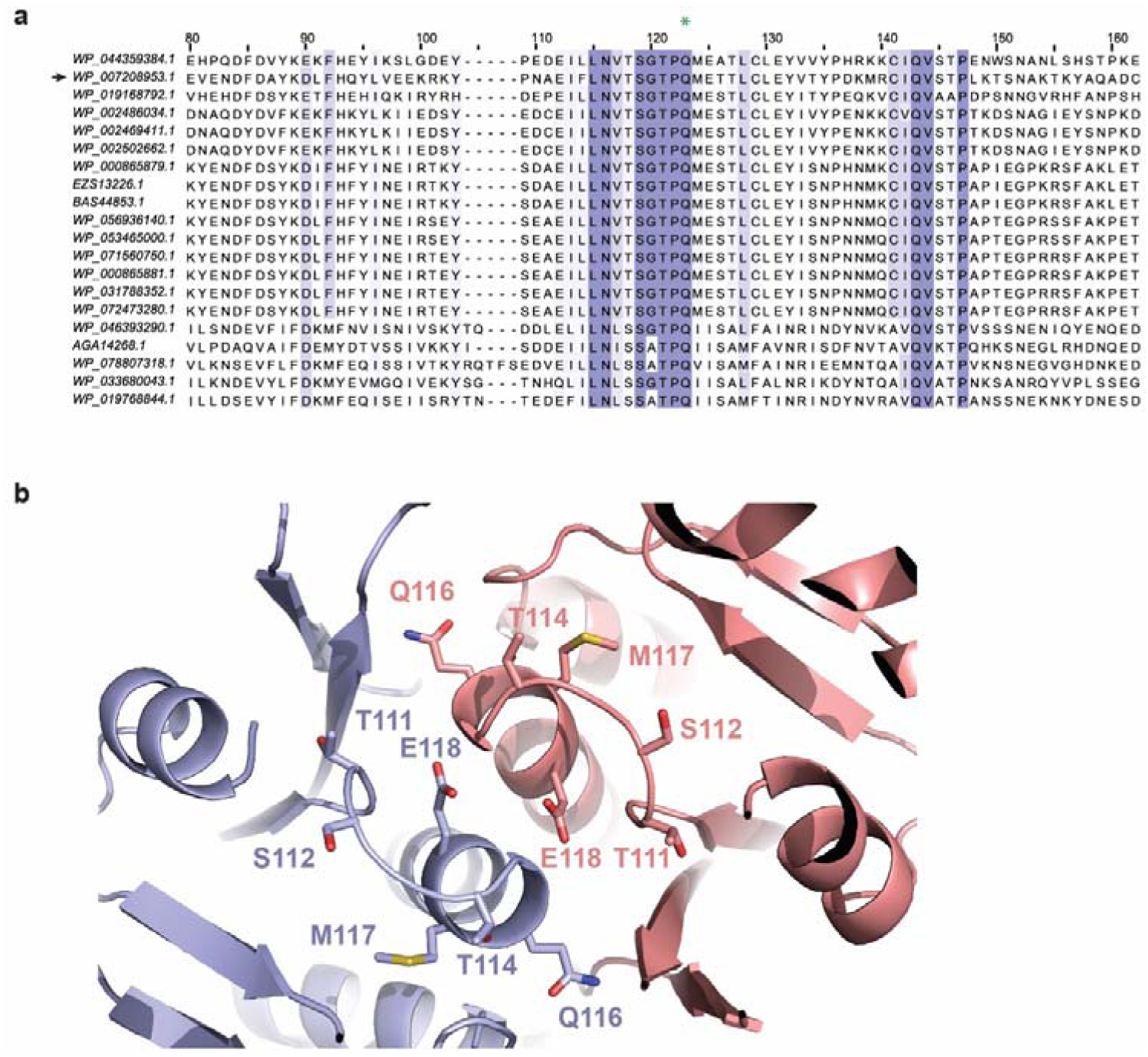
Conservation and structure of the CARF domain motif. a, Multiple sequence alignment of the CARF domain motifs in Csm6. The polypeptide sequence of EiCsm6 (denoted with an arrow) was used for a BLAST search against the NCBI non-redundant protein sequences database and 19 best scoring hits with >95 % query coverage were selected. The multiple sequence alignment was performed using Clustal Omega^33^ and visualized using ESPript 3^34^. Green asterisk the conserved glutamine residue mutated to abrogate allosteric activation of EiCsm6. **b,** Zoom-in view of a structural model of the dimeric interface of the CARF domains in the EiCsm6 dimer. The model was generated using Phyre2^35^.

**Extended Data Figure 5.**
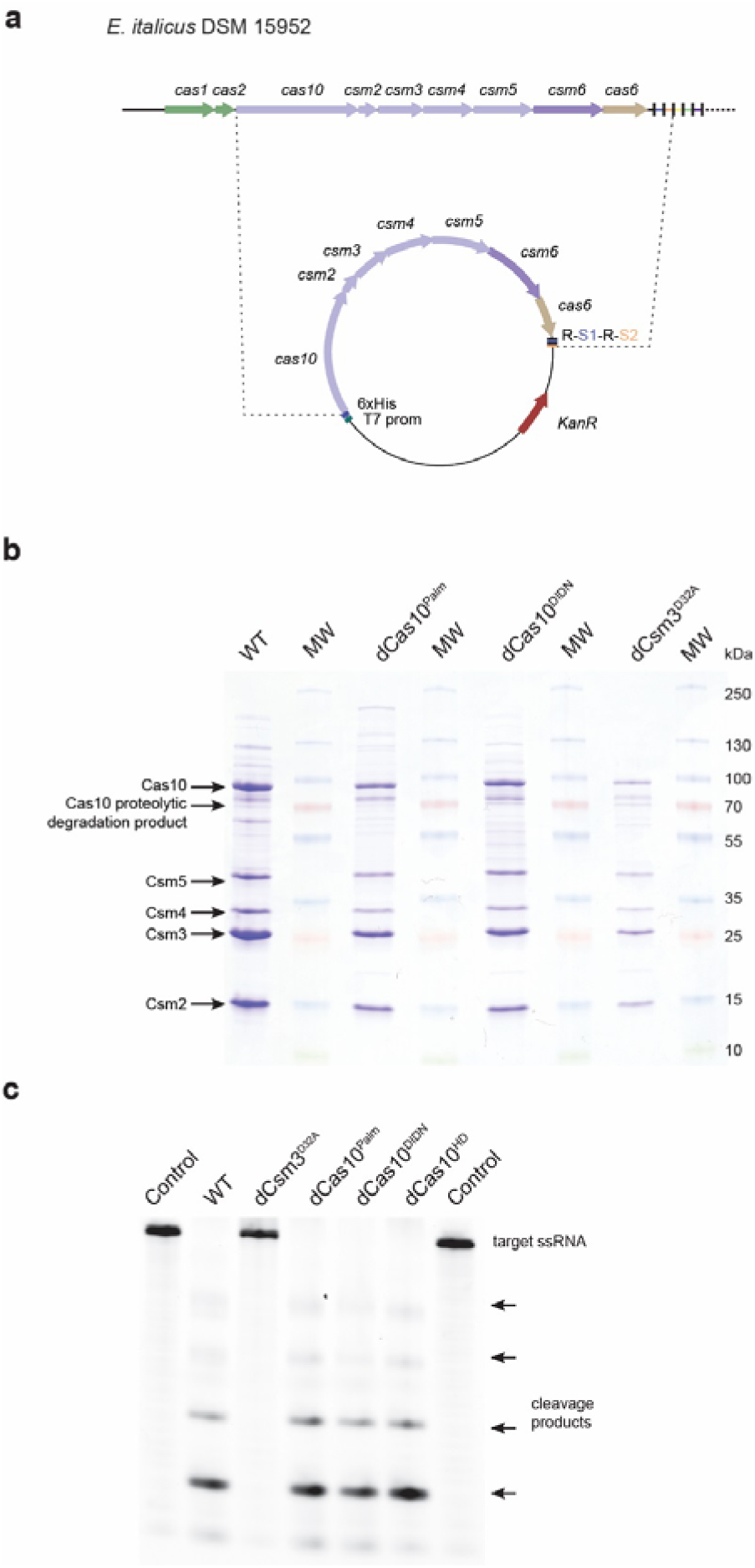
Purfication and crRNA-guided RNA cleavage by the EiCsm(1-5) complex. **a,** *Enterococcus italicus* DSM15952 contains a CRISPR-Cas locus with 9 *cas* genes followed by a repeat-spacer array. For heterologous expression of the EiCsm(1-5) complex, the genomic DNA fragment spanning genes *csm1*-*csm6* and *cas6* and the first two repeat-spacer units in the CRISPR array was cloned into the expression plasmid. **b,** SDSPAGE analysis of purified wild-type and mutant EiCsm(1-5) complexes. **c,** Denaturing gel of the products of a target RNase cleavage assay performed using a fluorophore-labelled target ssRNA substrate in the presence of WT or mutant EiCsm(1-5) complexes.

**Extended Data Figure 6.**
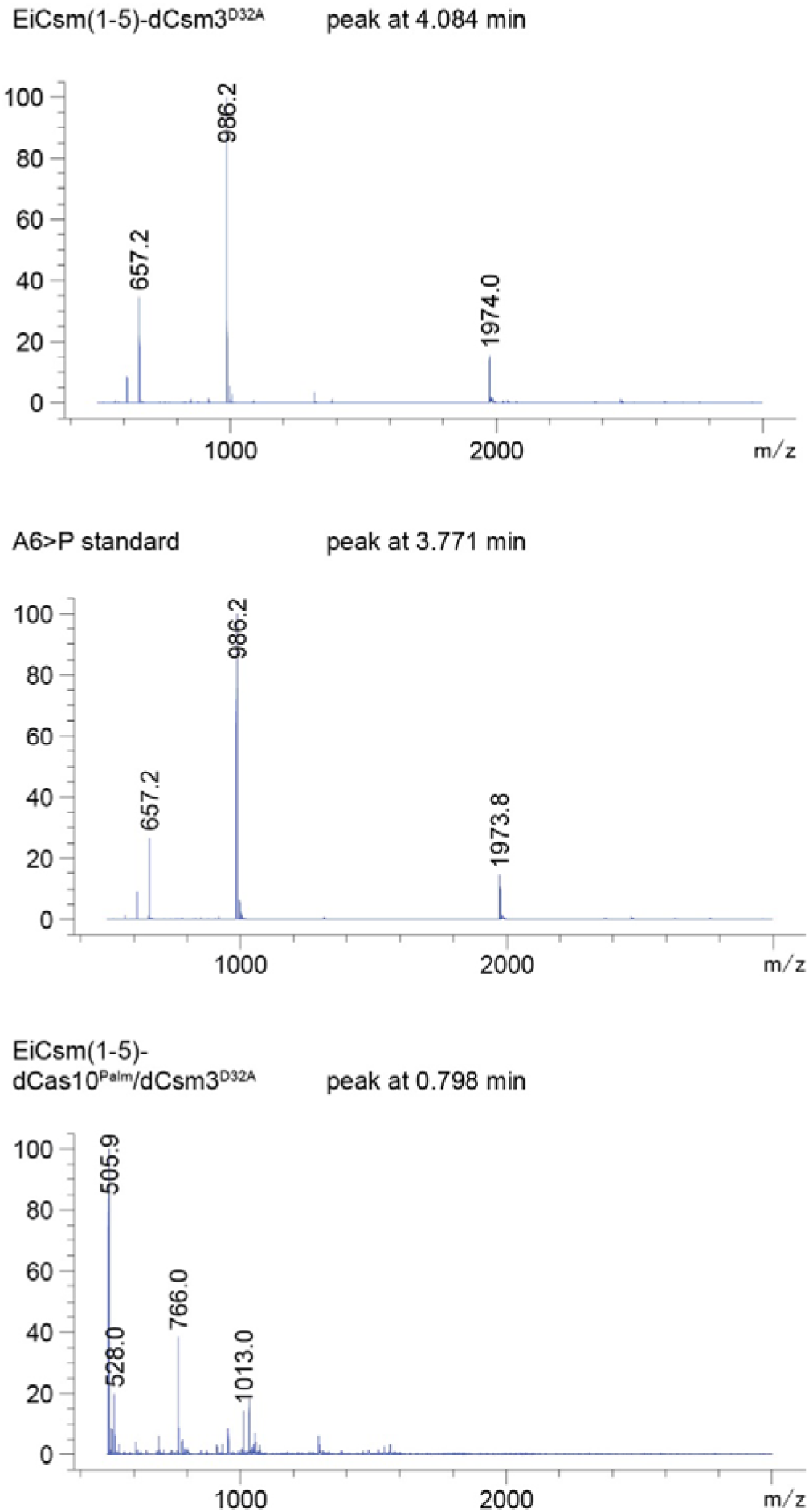
Mass spectrometry analysis of the Csm6 activator produced by the EiCsm(1-5) complex. Mass spectra of LC-MS peaks from the analysis of the products generated by the EiCsm(1-5)-dCsm3^D32A^ and EiCsm(1-5)-dCas10^Palm^/dCsm3^D32A^ complexes shown in Fig. 4c. The mass spectrum profile of synthetic A6>P standard is shown for comparison.

**Extended Data Figure 7.**
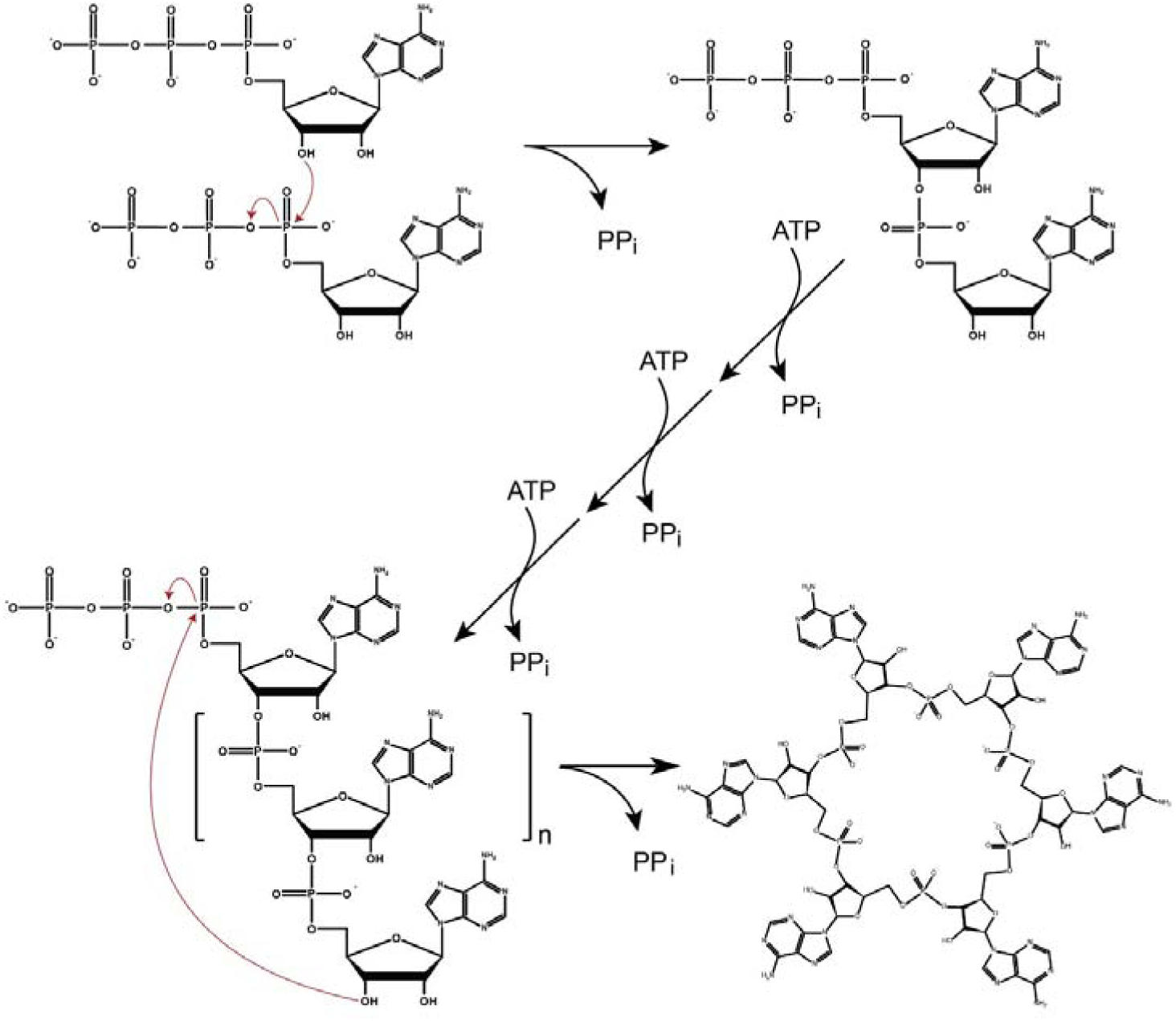
Proposed mechanism of cyclic oligoadenylate formation. The Cas10 Palm domain initially catalyses 5’-3’ phosphodiester bond formation by a nucleophilic attack of the 3’-hydroxyl group of one ATP substrate molecule onto the α-phosphate group of a second ATP molecule, yielding a 5’-triphosphorylated dinucleotide and inorganic pyrophosphate (PPi). The dinucleotide is extended by addition of successive AMP moieties to the 3’ end to generate a linear oligonucleotide. Final cyclisation involves the 3’-hydroxyl group attacking the α-phosphate group of the 5’-triphosphate moiety in the linear oligoadenylate intermediate. Notably, all reaction steps utilize the same catalytic mechanism and thus could be carried out by a single active site in the Palm domain of Cas10.

## REFERENCES

1. Mohanraju, P. et al. Diverse evolutionary roots and mechanistic variations of the CRISPR-Cas systems. Science 353, aad5147 (2016).

2. Marraffini, L. A. CRISPR-Cas immunity in prokaryotes. Nature 526, 55–61 (2015).

3. Sorek, R., Lawrence, C. M. & Wiedenheft, B. CRISPR-mediated adaptive immune systems in bacteria and archaea. Annu Rev Biochem 82, 237–266 (2013).

4. Koonin, E. V., Makarova, K. S. & Zhang, F. Diversity, classification and evolution of CRISPR-Cas systems. Curr Opin Microbiol 37, 67–78 (2017).

5. Makarova, K. S. et al. An updated evolutionary classification of CRISPR–Cas systems. Nat Rev Microbiol (2015). doi:10.1038/nrm icro3569

6. Samai, P. et al. Co-transcriptional DNA and RNA Cleavage during Type III CRISPR-Cas Immunity. Cell 161, 1164–1174 (2015).

7. Goldberg, G. W., Jiang, W., Bikard, D. & Marraffini, L. A. Conditional tolerance of temperate phages via transcription-dependent CRISPR-Cas targeting. Nature 514, 633–637 (2014).

8. Deng, L., Garrett, R. A., Shah, S. A., Peng, X. & She, Q. A novel interference mechanism by a type IIIB CRISPR-Cmr module in Sulfolobus. Mol. Microbiol. 87, 1088–1099 (2013).

9. Elmore, J. R. et al. Bipartite recognition of target RNAs activates DNA cleavage by the Type III-B CRISPR-Cas system. Genes Dev 30, 447–459 (2016).

10. Estrella, M. A., Kuo, F.-T. & Bailey, S. RNA-activated DNA cleavage by the Type III-B CRISPR-Cas effector complex. Genes Dev 30, 460–470 (2016).

11. Kazlauskiene, M., Tamulaitis, G., Kostiuk, G., Venclovas, Č. & Siksnys, V. Spatiotemporal Control of Type III-A CRISPR-Cas Immunity: Coupling DNA Degradation with the Target RNA Recognition. Mol Cell 62, 295–306 (2016).

12. Tamulaitis, G., Venclovas, Č. & Siksnys, V. Type III CRISPR-Cas Immunity: Major Differences Brushed Aside. Trends Microbiol. 25, 49–61 (2017).

13. Makarova, K. S., Anantharaman, V., Grishin, N. V., Koonin, E. V. & Aravind, L. CARF and WYL domains: ligand-binding regulators of prokaryotic defense systems. Front Genet 5, 102 (2014).

14. Anantharaman, V., Makarova, K. S., Burroughs, A. M., Koonin, E. V. & Aravind, L. Comprehensive analysis of the HEPN superfamily: identification of novel roles in intragenomic conflicts, defense, pathogenesis and RNA processing. Biol Direct 8, 15 (2013).

15. Hatoum-Aslan, A., Maniv, I., Samai, P. & Marraffini, L. A. Genetic characterization of antiplasmid immunity through a type III-A CRISPR-Cas system. J. Bacteriol. 196, 310–317 (2014).

16. Jiang, W., Samai, P. & Marraffini, L. A. Degradation of Phage Transcripts by CRISPR-Associated RNases Enables Type III CRISPR-Cas Immunity. Cell 164, 710–721 (2016).

17. Niewoehner, O. & Jinek, M. Structural basis for the endoribonuclease activity of the type III-A CRISPR-associated protein Csm6. RNA 22, 318–329 (2016).

18. Sheppard, N. F., Glover, C. V. C., Terns, R. M. & Terns, M. P. The CRISPR-associated Csx1 protein of Pyrococcus furiosus is an adenosine-specific endoribonuclease. RNA 22, 216–224 (2016).

19. Staals, R. H. J. et al. RNA targeting by the type III-A CRISPR-Cas Csm complex of Thermus thermophilus. Mol Cell 56, 518–530 (2014).

20. Tamulaitis, G. et al. Programmable RNA shredding by the type III-A CRISPR-Cas system of Streptococcus thermophilus. Mol Cell 56, 506–517 (2014).

21. Hatoum-Aslan, A., Samai, P., Maniv, I., Jiang, W. & Marraffini, L. A. A Ruler Protein in a Complex for Antiviral Defense Determines the Length of Small Interfering CRISPR RNAs. Journal of Biological Chemistry 288, 27888–27897 (2013).

22. Corrigan, R. M. & Gründling, A. Cyclic di-AMP: another second messenger enters the fray. Nat Rev Microbiol 11, 513–524 (2013).

23. Gaffney, B. L., Veliath, E., Zhao, J. & Jones, R. A. One-flask syntheses of c-di-GMP and the [Rp,Rp] and [Rp,Sp] thiophosphate *analogues*. Org. Lett. 12, 3269–3271 (2010).

24. Carroll, S. S. et al. Activation of RNase L by 2’,5’-oligoadenylates. Kinetic characterization. J Biol Chem 272, 19193–19198 (1997).

25. Prischi, F., Nowak, P. R., Carrara, M. & Ali, M. M. U. Phosphoregulation of Ire1 RNase splicing activity. Nat Commun 5, 3554 (2014).

26. Staals, R. H. J. et al. Structure and Activity of the RNA-Targeting Type III-B CRISPRCas Complex of Thermus thermophilus. Mol Cell 52, 135–145 (2013).

27. Makarova, K. S. et al. Evolution and classification of the CRISPR-Cas systems. Nat Rev Microbiol 9, 467–477 (2011).

28. Osawa, T., Inanaga, H. & Numata, T. Crystal Structure of the Cmr2-Cmr3 Subcomplex in the CRISPR-Cas RNA Silencing Effector Complex. J Mol Biol 425, 3811–3823 (2013).

29. Yan, X., Guo, W. & Yuan, Y. A. Crystal structures of CRISPR-associated Csx3 reveal a manganese-dependent deadenylation exoribonuclease. RNA biology 12, 749–760 (2015).

30. Kalia, D. et al. Nucleotide, c-di-GMP, c-di-AMP, cGMP, cAMP, (p)ppGpp signaling in bacteria and implications in pathogenesis. Chem Soc Rev 42, 305–341 (2013).

31. Hornung, V., Hartmann, R., Ablasser, A. & Hopfner, K.-P. OAS proteins and cGAS: unifying concepts in sensing and responding to cytosolic nucleic acids. Nat. Rev. Immunol. 14, 521–528 (2014).

32. Lintner, N. G. et al. The structure of the CRISPR-associated protein Csa3 provides insight into the regulation of the CRISPR/Cas system. J Mol Biol 405, 939–955 (2011).

33. Sievers, F. et al. Fast, scalable generation of high-quality protein multiple sequence alignments using Clustal Omega. Molecular Systems Biology 7, 539–539 (2011).

34. Gouet, P., Courcelle, E., Stuart, D. I. & Métoz, F. ESPript: analysis of multiple sequence alignments in PostScript. Bioinformatics 15, 305–308 (1999).

35. Kelley, L. A., Mezulis, S., Yates, C. M., Wass, M. N. & Sternberg, M. J. E. The Phyre2 web portal for protein modeling, prediction and analysis. Nat Protoc 10, 845–858 (2015).

36. Geertsma, E. R. FX cloning: a simple and robust high-throughput cloning method for protein expression. Methods Mol. Biol. 1116, 153–164 (2014).

37. Kreiswirth, B. N. et al. The toxic shock syndrome exotoxin structural gene is not detectably transmitted by a prophage. Nature 305, 709–712 (1983).

